# Active control of mitochondrial network morphology by metabolism driven redox state

**DOI:** 10.1101/2024.08.05.606562

**Authors:** Gaurav Singh, Vineeth Vengayil, Aayushee Khanna, Swagata Adhikary, Sunil Laxman

## Abstract

Mitochondria are dynamic organelles that constantly change morphology. What controls mitochondrial morphology however remains unresolved. Using actively respiring yeast cells growing in distinct carbon sources, we find that mitochondrial morphology and activity are unrelated. Cells can exhibit fragmented or networked mitochondrial morphology in different nutrient environments independent of mitochondrial activity. Instead, mitochondrial morphology is controlled by the intracellular redox state, which itself depends on the nature of electron entry into the Electron Transport Chain (ETC)— through complex I/II, or directly to coenzyme Q/cytochrome c. In metabolic conditions where direct electron entry is high, reactive oxygen species (ROS) increase, resulting in an oxidized cytosolic environment and rapid mitochondrial fragmentation. Decreasing direct electron entry into the ETC genetically or chemically, or reducing the cytosolic environment rapidly restores networked morphologies. Using controlled disruptions of electron flow to alter ROS and redox state, we demonstrate minute-scale, reversible control between networked and fragmented forms in an activity independent manner. Mechanistically, the fission machinery through Dnm1 responds in minute-scale to redox state changes, preceding the change in mitochondrial form. Thus, the metabolic state of the cell and its consequent cellular redox state actively controls mitochondrial form.

## Introduction

Mitochondria are double-membrane bound organelles at the core of energetics and cell state. They play a central role in ATP production through the functioning of the electron transport chain (ETC) and oxidative phosphorylation, coupled with the tricarboxylic acid (TCA) cycle - that also produces precursors for the synthesis of lipids, amino acids, and nucleotides (1–3). The TCA cycle supplies electron donors (NADH and FADH2) to ETC. Electrons flow through the enzyme complexes of the ETC, and generate the proton-motive force across the inner mitochondrial membrane that drives the ATP synthase for ATP generation. Even as the mitochondria carries out electron transfer in the context of ATP generation, the process of electron transport through the respiratory chain is not completely leak-proof and results in generating reactive oxygen species (ROS), a byproduct of respiration (4, 5). Far from being static biochemical reactors for energetics, mitochondria are dynamic organelles (6) that exhibit a striking diversity in form, ranging from a connected network of tubules to round, isolated fragments under different environments, or during cell cycle progression (7). Mitochondria undergo remodeling by fission and fusion to maintain their form. These processes are thought to allow mitochondria to adapt to various environments, or ensure optimal energy production (8, 9). Indeed, how mitochondrial networks respond to diverse environments have attracted intense scientific inquiry, since mitochondrial dysfunction is implicated in a wide range of diseases (10, 11).

However, we lack an empirical understanding of a relationship between mitochondrial form and function, or indeed what controls mitochondrial form. Assumptions have been made regarding the association of mitochondrial morphology with mitochondrial activity (12–14)). For example, associations have been made with interconnected mitochondrial networks and high respiratory activity, or fragmented mitochondria that have been observed in quiescent or inactive cells (10). There are numerous contrasting examples where mitochondrial form does not go hand in hand with activity. For example, Romani et al (15) find drastic changes in mitochondrial morphology without changes in respiration, or Yu et (16) find conditions where there is mitochondrial fragmentation without changes in activity. In stem cells, different mitochondrial morphologies have been observed, with unclear relationships with bioenergetics or activity (17–19). Therefore, the underlying basis for what triggers mitochondrial network remodeling remains unresolved.

In contrast, we understand the molecular machinery that regulates mitochondrial form. Mitochondrial morphology is regulated by a balance of fission and fusion processes, through specific proteins and mechanisms that are well conserved across eukaryotic cells, ranging from yeast to mammals (20). Mitochondrial fission is spearheaded by large dynamin family GTPase, Drp1 (Dnm1 in yeast) that is recruited on mitochondrial membrane by accessory proteins (Mff, Mid49/51 and Fis1), and act in unison with ER and actin-myosin cytoskeleton to divide mitochondria (21). The fusion of outer and inner-mitochondrial membranes is facilitated by mitofusins, Mfn1 and Mfn2, (Fzo1 in yeast) and Opa1 (Mdm1 in yeast). The fission and fusion machinery are heavily regulated (22), but have predominantly been studied in the context of extreme starvation or mitochondrial damage, where mitochondrial quality control systems ensure hyper-fused mitochondria (23), or fragmented mitochondria (24). In summary, the regulators of the fission and fusion process in mitochondria (25) are well studied. In contrast, the initiating signals (26) that determine mitochondrial form remain obscure.

In this study, we address what controls mitochondrial form by examining budding yeast (*S. cerevisiae*) grown in multiple defined carbon-sources, where the respective requirements for mitochondrial function and respiration are established (27). We quantitatively estimate the extent and nature of the mitochondrial networks in these cells, while assessing multiple indicators of mitochondrial outputs. Through this we find that mitochondrial form depends on the collective redox state, and is independent of activity. Specific metabolic conditions result in increased direct electron entry via coenzyme Q or cytochrome c into the ETC, which consequently increases ROS. This ROS directly results in fragmented mitochondria. Reducing direct electron entry (through genetic approaches) results in networked mitochondria, as does decreasing ROS through chemical approaches. Orthogonally increasing ROS due to oxidants rapidly fragments mitochondria. In contrast, a reduced cytosolic environment - which is a consequence of the metabolic state of the cell - leads to networked mitochondria. This control of mitochondrial form is rapid (within minutes), dynamic, fully reversible, and independent of mitochondrial activity. Finally, the activity of the fission-regulating protein Dnm1 rapidly responds to changes in redox state, to control mitochondrial morphology. Overall, our data suggest a systems-level chemical framework for how distinct metabolic activities controls cytosolic redox state, to regulate mitochondrial form via the fission machinery, and independent of mitochondrial activity. Our study opens direct lines of inquiry on the nature of redox signals that control mitochondrial form via the activity of the fission machinery.

## Results

### Mitochondrial network morphology does not correlate with activity

Mitochondria have morphologies that range from punctate, small and fragmented to interconnected tubules, that are observed in different cells and conditions (28, 6, 29, 8, 30). These mitochondrial forms dynamically change in response to environmental factors (8, 12, 13, 15, 31, 32). It is currently unclear if mitochondrial form is related to mitochondrial function (Fig. 1A). In part, any correlations made are difficult to generalize, because observations made across different cell/tissue types or organisms cannot be directly compared. A practical lacuna has been the absence of studies using precisely manipulable systems where mitochondrial morphology can be predictably varied in controlled, experimental settings. The budding yeast (*Saccharomyces cerevisiae*), has been an ideal model system to identify general principles of mitochondrial function (33–38). Yeast cells are metabolically versatile, and grow on different carbon sources with varying requirements of mitochondrial respiration. In glucose, yeast repress their mitochondrial outputs via the Crabtree effect (39). In other carbon sources, yeast cells have higher requirements for mitochondrial respiration to sustain cellular energetics. Therefore, using yeast growing in different carbon sources, it would be possible to directly compare mitochondrial activity and morphology, in precisely defined settings.

**Fig. 1.**
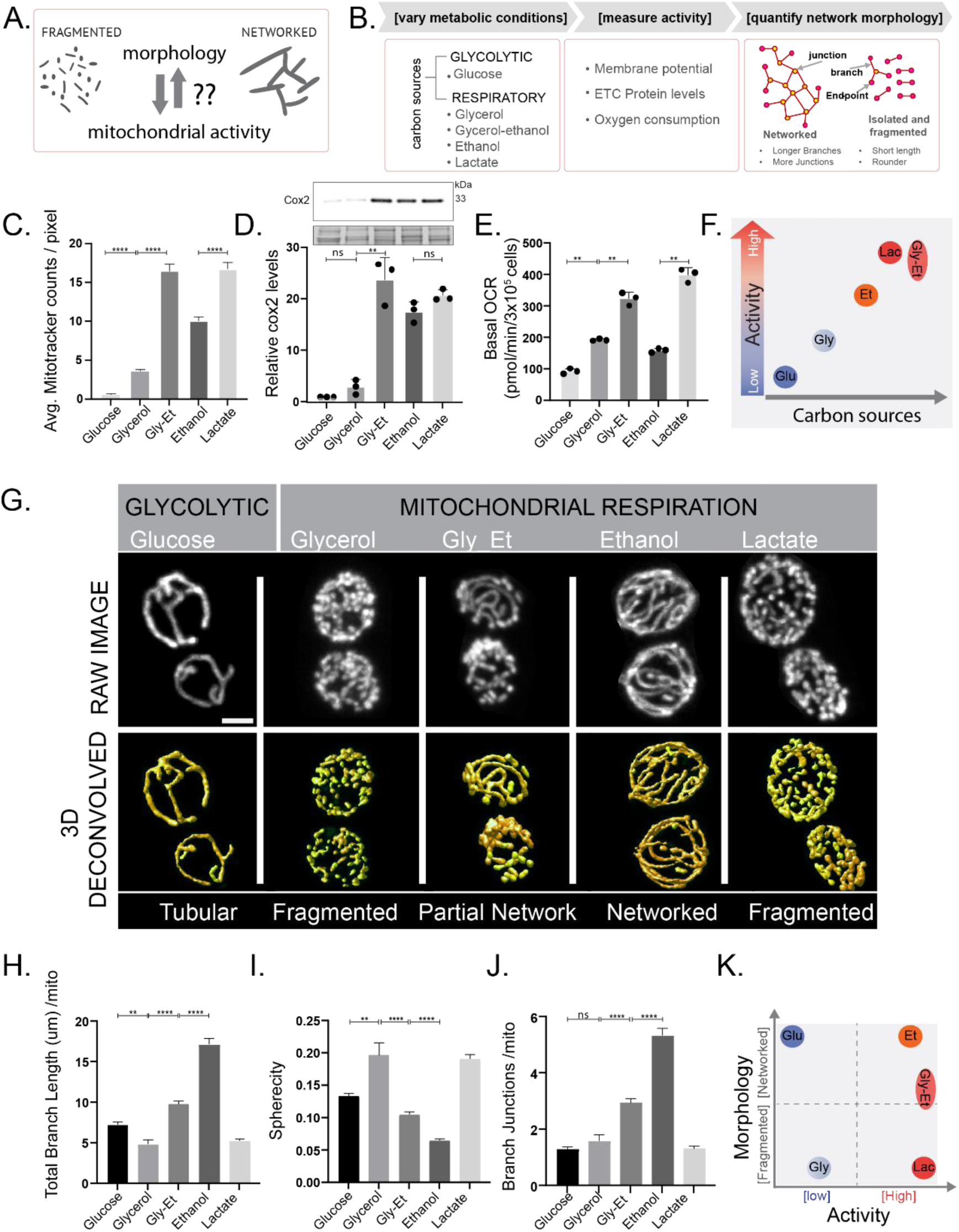
Mitochondrial morphology does not correlate with activity in different nutrient environments. (A) Schematic asking if mitochondrial network morphologies and activities are interconnected. (B) Experiment design to investigate this relationship that involves growing cells in five different carbon sources as indicated, measuring mitochondrial activity using standardized assays (C-E), and quantifying mitochondrial morphology (G-J). Measurement of multiple measures of mitochondrial activity in different carbon sources:(C) Average Mitotracker fluorescence that reports on mitochondrial membrane potential, measured by single cell imaging. Data represents average intensity per pixel for >100 cells from three independent experiments (mean ±SEM). (D) ETC complex IV subunit Cox2 protein levels measured by western blot using an anti-Cox2 antibody. A representative blot (out of 3 biological replicates, n=3) and their quantifications are shown, and (E) Basal oxygen consumption rate (OCR) corresponding to ∼3 x 10^5 cells, from three independent experiments (n=3), normalized to the OD600. (F) Schematic summary of mitochondrial activity of cells grown in different carbon sources. All respiratory carbon sources (glycerol, glycerol ethanol, ethanol and lactate) show higher baseline mitochondrial activity as compared to cells grown in glucose. However, there is considerable heterogeneity in morphology and mitochondrial activities even amongst various respiratory carbon sources. (G) Representative microscopy images of cells with mitochondria targeted with mNeonGreen. Top panel shows the maximum intensity projection of raw 3D z-stack and bottom panel shows 3D renderings of the z-stack after deconvolution. (H-J) Quantitative 3D analysis and comparison of mitochondrial network morphology across different carbon sources. Data represents analysis of more than 200 cells from three independent experiments (mean ±SEM). Scale bar - 2µm. (K) Schematic summary of mitochondrial activity versus respective mitochondrial morphology for cells grown in different carbon sources. Data represents mean ± SD unless otherwise stated. *P < 0.05, **P < 0.01, and ***P< 0.001; n.s., non-significant difference, calculated using unpaired Student’s *t* tests.

We first established an experimental and analytical pipeline where cells were grown in a defined carbon source, where we could quantitatively assess mitochondrial morphology as well as activity (Fig 1B). The carbon sources used were glucose (repressed mitochondrial respiration), ethanol, glycerol, glycerol-ethanol, and lactic acid (all of which require mitochondrial respiration). We first measured multiple parameters of mitochondrial activity - mitochondrial membrane potential, oxygen consumption, and amounts of a key respiratory complex IV protein Cox2 (Fig 1B), to obtain a composite assessment of activity. We separately obtained high-resolution images of mitochondria, and estimated various network parameters. For accurate imaging and quantification of mitochondrial network, 3D confocal z-stacks were acquired that were then deconvolved. The 3D deconvolved stack is further processed for 3D multidimensional analysis of mitochondrial morphology in ImageJ. Supplementary methods and Supplementary Fig S1A-C provide detailed information on the imaging pipeline used, and addresses critical aspects of image deconvolution and 3D image processing to obtain clear separation of individual mitochondria, and accurate reconstructions of the mitochondrial networks.

Mitochondrial activity in yeast cells depends on the carbon source available. In glucose, cells repress mitochondrial activity (39). In sole carbon sources such as ethanol, lactate or glycerol, the repression due to glucose is released, and cells carry out mitochondrial respiration. We first estimated mitochondrial membrane potential in cells under all these conditions using a rosamine based potentiometric dye - Mitotracker red (39, 40). Expectedly, mitochondrial potential was low in glucose (Fig 1C). However, mitochondrial potential showed different extents of increase in glycerol, ethanol, glycerol-ethanol, or lactate (Fig 1C), indicating distinct membrane potentials. We next estimated relative amounts of Cox2 protein, as an indicator of the ETC (39, 41). Glucose had low Cox2 amounts, while a clear hierarchy of increased Cox2 protein was observed in the other carbon sources (Fig 1D). Finally, we measured basal oxygen consumption (using a Seahorse based assay (39, 42)) in cells in these conditions (Fig 1E). Consistent with the other observations, cells in glucose had very low oxygen consumption, while in each of the other conditions, oxygen consumption was high, but with a clear hierarchy (Fig 1E). Collectively, cells grown in lactate and glycerol-ethanol show the highest levels of mitochondrial activity, cells in ethanol have high activity, and cells growing in glycerol have relatively low activity, as summarized in Fig 1F.

We next asked what is the nature of mitochondrial networks in these conditions. Fig 1G shows the observed mitochondrial morphologies in each of these different carbon sources. Cells in glucose have few but well-connected mitochondrial tubules. In contrast, cells grown in different respiratory carbon sources showed distinct mitochondrial networks (Fig 1G). The morphologies group into - 1) Completely fragmented mitochondria, characterized by short and round tubules as observed in glycerol and lactate (also see Supplementary Video S1), 2) Well networked mitochondria with long and interconnected mitochondrial tubules as observed in ethanol (see Supplementary Video S1), and 3) Partially networked mitochondria with long connected tubules intermixed with shorter tubules as seen in glycerol-ethanol (Fig 1G). We carried out an unbiased, automated 3D quantification of mitochondrial network (Fig 1H-J) in all the carbon sources, to quantify the visual morphological features based on average branch length, number of branches and branch junctions per mitochondria, average mitochondrial sphericity and form factor (Fig 1H-J, Supplementary Fig S1D and S1E, and see Material and Methods for details).

In summary, there is no correlation of mitochondrial morphology with mitochondrial activity. For example, cells grown in glycerol or lactate both show highly fragmented mitochondria but have very low or very high mitochondrial activities respectively. Contrastingly, cells grown in ethanol and glucose both have long, highly networked mitochondrial tubules but high or low mitochondrial activities respectively. Cells in glucose or glycerol have low activities, but networked or fragmented mitochondria respectively. Therefore, in cells growing in a range of commonly occurring growth environments, mitochondrial morphology does not correlate with activity (Fig 1K and S1F).

### Cellular redox state correlates with mitochondrial morphology

Since mitochondrial morphology did not correlate with activity, what might explain the distinct morphologies observed in different metabolic conditions? To ask this question, we wondered if other mitochondrial outputs might be relevant. It is well established that mitochondrial outputs contribute to changing the overall cellular redox state, via the production of reactive oxygen species (ROS) that diffuse out of mitochondria (43–45). Interestingly, it is not as clear under what contexts does mitochondrial activity result in increased ROS. We therefore wondered whether cells growing in distinct carbon sources (where mitochondrial morphologies are very different) had different intracellular redox environments (Fig. 2A). In order to monitor differences in intracellular oxidation levels of cells grown in distinct carbon sources, we first used a well-established dye-based fluorescence assay (DCFDA fluorescence) as a quantitative readout. DCFDA is a cell-permeable non-fluorescent dye that becomes fluorescent upon oxidation, and is used to measure differences in intracellular redox states (15, 16, 46). Cells grown in glycerol and lactate show high DCFDA fluorescence - consistent with a more oxidized intracellular environment - as compared to cells grown in ethanol, ethanol-glycerol or glucose (Fig 2B). This data suggested that cells in glycerol and lactate have a more oxidized cellular state. Contrastingly, cells grown in ethanol, glycerol-ethanol or glucose all exhibited lower DCFDA fluorescence - which would be consistent with a more reduced intracellular environment (Fig 2B). As a control, we supplemented ethanol-grown cells with hydrogen-peroxide (an oxidizing agent), which led to an increase in DCFDA fluorescence (Fig 2B).

**Fig. 2.**
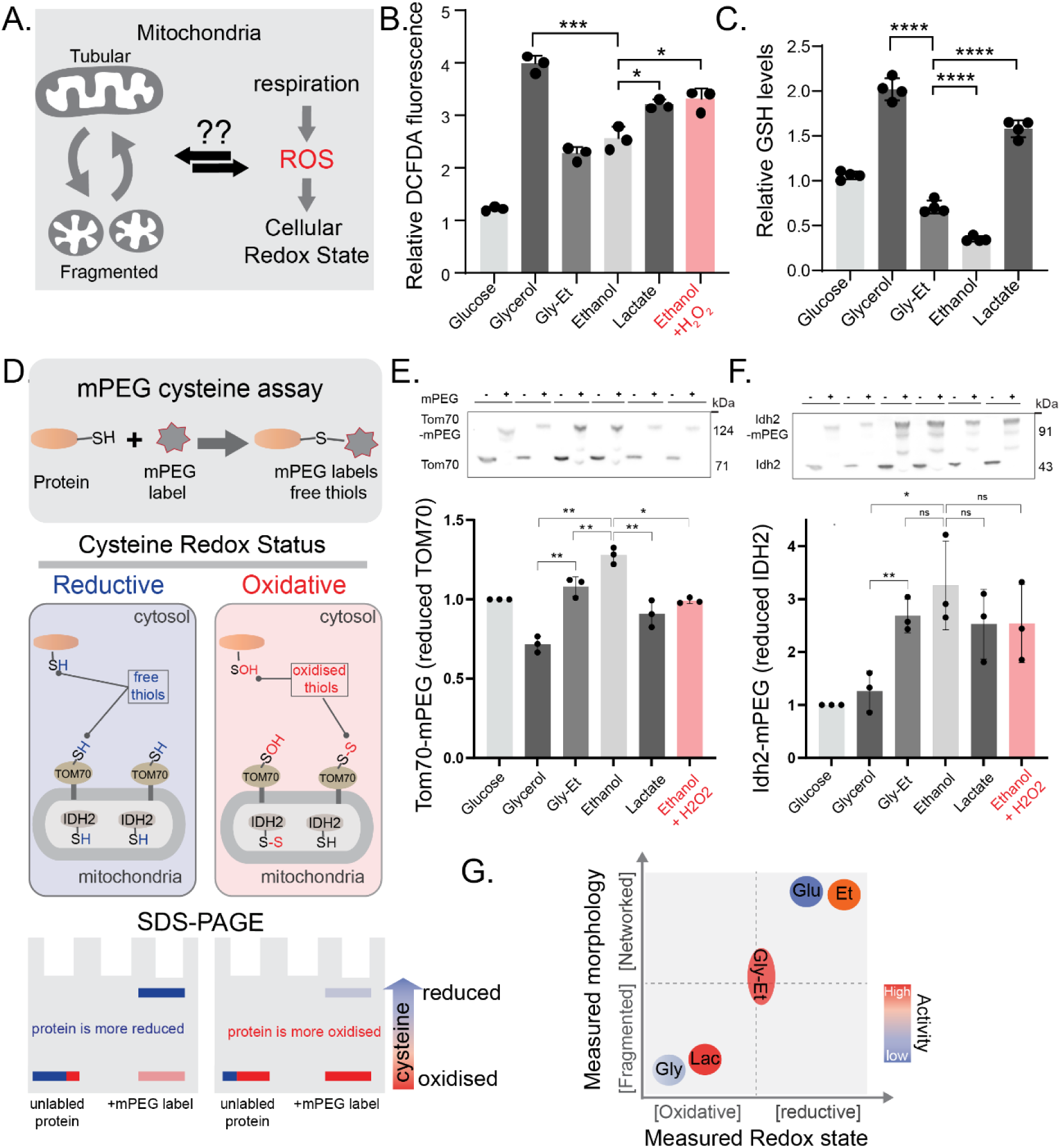
Relationship of cellular oxidative state and mitochondrial morphology. (A) Schematic showing whether respiration dependent changes in cellular redox state can explain changes in mitochondrial morphology. (B) The total cellular redox state (measured by DCFDA fluorescence) in cells grown in different carbon sources. Cells were grown in different media as described before and the relative DCFDA fluorescence was measured (see materials and methods). (C) The relative total cellular GSH levels in cells grown in different carbon sources. Cells were grown in different media as described and GSH levels were measured by LC-MS/MS. (D) Schematic showing maleimide-PEG (mPEG) assay for protein redox state estimation. (E) Cytosolic oxidative state in different carbon sources. Cells were grown in different media as described and oxidation state of the cytosol was measured by electrophoretic mobility shift of mPEG bound to cytosolic facing Tom70, in a western blot (using anti-FLAG antibody). (F) Mitochondrial oxidative state in different carbon sources. Cells were grown in different media as described and oxidation state of mitochondria was measured by electrophoretic mobility shift of mPEG bound to mitochondrial Idh2, in a western blot (using anti-FLAG antibody). Data represents mean ± SD from three biological replicates (n=3) for all the experiments. (G) Schematic summary of how mitochondrial morphology correlates with cellular redox state in cells grown in different carbon sources with varied mitochondrial activities. *P < 0.05, **P < 0.01, and ***P< 0.001; n.s., non-significant difference, calculated using unpaired Student’s *t* tests for data in (A) and (B) and paired Student’s *t* tests for data in (E) and (F).

Oxidation states cannot be interpreted based solely on DCFDA fluorescence. Therefore, in order to have an alternate readout of cellular oxidation levels, we next measured relative amounts of total glutathione (GSH) in cells in each of these conditions using mass-spectrometry. GSH is a primary redox buffer in cells, functioning as an antioxidant against ROS. Therefore, higher glutathione amounts are expected in more oxidizing conditions, as cells counter this environment (15, 47, 48). Notably, we observed significantly higher GSH amounts in lactate and glycerol (Fig 2C) as compared to cells growing in ethanol, ethanol-glycerol, or glucose – consistent with cells experience a more oxidizing environment in lactate and glycerol, and increasing GSH in response.

However, DCFDA fluorescence and total GSH amounts reflect gross intracellular redox environments. We therefore devised a biochemical assay to assess the relative oxidative state of the cytosolic or mitochondrial environment in the different carbon sources. For this, we utilized a maleimide-PEG (49) (polyethylene glycol) based assay (50) to biochemically measure the relative oxidation status of cysteine residues present on select proteins which are located either inside the mitochondrial matrix (isocitrate dehydrogenase - Idh2), or on the mitochondrial surface facing/exposed to the cytosol (Tom70), or cytosolic GAPDH. The maleimide-PEG will covalently conjugate to reduced (and not oxidized) cysteine residues, and result in an added mass on the protein that is visualized via mobility changes on a denaturing SDS-PAGE gel. This assay and its expected outputs are illustrated in the schematic diagram (Fig 2D). Through these experiments, we observed that the cytosol facing Tom70 had more reduced cysteines (i.e. more cysteines that can react with the added maleimide-PEG) in cells grown in ethanol or ethanol-glycerol, compared to cells grown in lactate or glycerol (Fig 2E). In contrast, Idh2 (which is present inside the mitochondrial matrix) showed no significant changes in cysteine oxidation across ethanol, ethanol-glycerol, or lactate (Fig 2F). Finally, cytosolic GAPDH showed more reduced cysteines in cells grown in ethanol or ethanol-glycerol, compared to lactate or glycerol (Supplementary S2). Results of these direct assays estimating the relative oxidation status of the cytosolic environment suggest that cells growing in ethanol or glycerol-ethanol have a more reduced intracellular (cytosolic) environment, while cells in glycerol or lactate have a more oxidized environment.

With these observations, we can summarize the relationship of relative oxidative states with mitochondrial morphology (Fig 2G). The mitochondrial morphology in different metabolic conditions correlates with intracellular redox environment, independent of activity. In metabolic conditions where intracellular environment is more oxidized, mitochondria show fragmented morphology whereas metabolic conditions that have a relatively reduced intracellular environment exhibit more networked mitochondria.

### Mitochondrial morphology is reversibly and rapidly controlled by redox state

From our previous observations, the mitochondrial morphology correlated with the intracellular redox state, where a more oxidized intracellular environment had fragmented mitochondria. Therefore, we wondered if it is possible to alter mitochondrial morphology in a predictable fashion (Fig 3A), solely by manipulating the redox environment?

**Fig. 3.**
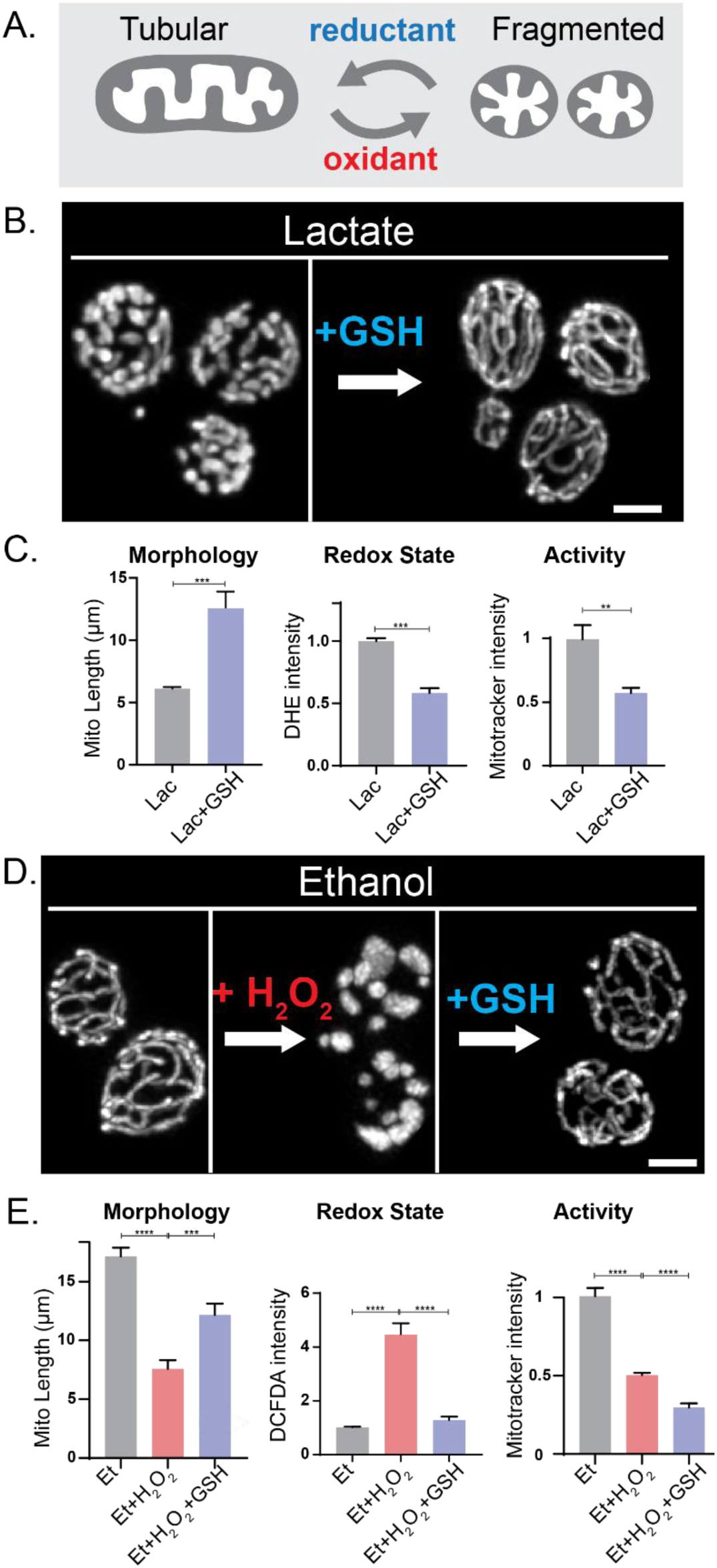
Mitochondrial network morphology changes in response to redox perturbations. (A) Schematic showing whether mitochondrial morphology can be reversibly modulated by changing redox state upon addition of oxidants or antioxidants. (B) Cells grown in lactate upon incubation with Glutathione (GSH) for 30 mins show a reversal in mitochondrial morphology from fragmented to networked (quantified in (C)). (D) Cells grown in ethanol are first incubated with H_2_O_2_ for 30 mins and then further incubated with GSH for 30 min. Addition of H_2_O_2_ fragments mitochondria that is then restored upon further addition of GSH. For data shown in panels (B) and (D), changes in mitochondrial morphology, redox state and activity are quantified in (C) and (E) respectively. Mitochondrial morphology is assessed by quantifying average mitochondrial length per mitochondria, redox state is measured by evaluating changes in DCFDA or DHE fluorescence and Mitotracker fluorescence is used as a proxy for measuring changes in activity. Data represents mean ±SEM (see material and methods for details). Scale bar - 2µm. *P < 0.05, **P < 0.01, and ***P< 0.001; n.s., non-significant difference, calculated using unpaired Student’s *t* tests.

To address this question, first we used cells growing in lactate, which is characterized by an oxidized state and fragmented mitochondria. Upon introducing an antioxidant - glutathione or ascorbic acid - to these cells growing in lactate, within ∼30 minutes we observed a rapid transition to a fully networked mitochondrial morphology (Fig 3B). In these cells we quantified morphology, intracellular redox state and mitochondrial potential (as a proxy for activity). Notably, upon the addition of the antioxidant, the mitochondria rapidly (within 30 min) became networked, the intracellular environment became more reduced, although the mitochondrial activity decreased (Fig 3C). In a complementary experiment, we cultured cells in ethanol (reduced intracellular environment, mitochondria are highly networked). Adding an oxidant (H_2_O_2_) to these cells caused their mitochondria to fragment within 30 minutes (Fig 3D). When we subsequently added an antioxidant (reduced glutathione or ascorbic acid) to these oxidant-treated cells, the mitochondrial network morphology was fully restored within 30 minutes (Fig 3D, Fig S3). We measured the same parameters: morphology, redox state, and activity. The oxidant induced mitochondrial fragmentation and an oxidized intracellular environment, which were both reversed upon addition of the antioxidant (Fig 3E). Notably, even while the mitochondrial morphology responded to changes in redox state, the mitochondrial activity consistently decreased and did not correlate with changes in morphology (Fig 3E).

Concluding, these data show that the mitochondrial network morphology in any given environment can be reversibly controlled by changing the intracellular redox environment. Mitochondrial networks fragment in a more oxidized environment and vice versa. These changes in morphology do not correlate with mitochondrial activity.

### Metabolic contexts resulting in direct electron entry have fragmented mitochondria

These observations now raise the question - in different nutrient environments (all of which require mitochondrial activity), what might be a reason for this altered oxidative state. Why might there be a more oxidized cytoplasm in some conditions, without a correlation with mitochondrial activity? We reasoned that one possibility was that there might be intrinsic differences in how each carbon source is metabolized by the mitochondria, that lead to changes in ROS levels. To interrogate this hypothesis, we first explored how each of these carbon sources is metabolized by yeast - and how the electron transport chain functions in these different contexts. In the schematic shown in Fig 4A, we summarize how each of these carbon sources are metabolized, with the ETC function placed in context. We can immediately observe contrasting processes of how each of these carbon sources drive the ETC (Fig 4A, Fig 4B). Ethanol is a classical ‘respiratory’ carbon source that yeast utilize after depleting glucose. When ethanol is metabolized, it enters the TCA cycle, from where the reducing equivalents (NADH/FADH2) are produced to drive the ETC (Fig 4A). In contrast, lactate is metabolized by a lactate dehydrogenase (LDH) enzyme first. In *S. cerevisiae*, the primary yeast lactate dehydrogenase - Cyb2 (27, 51, 52) (L-lactate cytochrome-c oxidoreductase) is unusual in that the electrons are not donated to NAD, but to ferricytochrome c, and thus this electron directly enters the respiratory chain via cytochrome c (Fig 4A). Similarly, glycerol is metabolized by the glycerol-3-phosphate dehydrogenase (GPDH enzyme)) (53, 54). The primary GPDH enzyme in yeast is Gut2, and this enzyme directly injects an electron into the ETC via a ubiquinone (Fig 4A). Therefore, both lactate and glycerol metabolism drive mitochondrial function via direct electron injection into the ETC, via a cytochrome or a ubiquinone (by bypassing complex 1 and/or 2). Both these enzymes are present on the outer leaflet of the inner-mitochondrial membrane. Furthermore, the binding site/pocket of these enzymes for redox acceptors (quinone or cytochrome c) might not be as well protected and shielded as compared to canonical ETC complex I and complex II where the quinone binding pocket is located much deeper. Therefore, we hypothesized that this direct injection of electrons into the ETC by either of these enzymes would result in more electrons interacting with oxygen, leading to higher ROS generation (Fig 4A and Fig 4B), as postulated previously (53, 55, 56). We can therefore observe that in ethanol or glucose (with tubular/networked mitochondria) the reducing equivalents produced via the TCA cycle drive the ETC by conventional electron transfer to complex I and complex II (Fig 4B). In contrast, in lactate or glycerol (fragmented mitochondria), there is direct electron injection into the ETC during the metabolism of these carbon sources (Fig 4B).

**Fig. 4.**
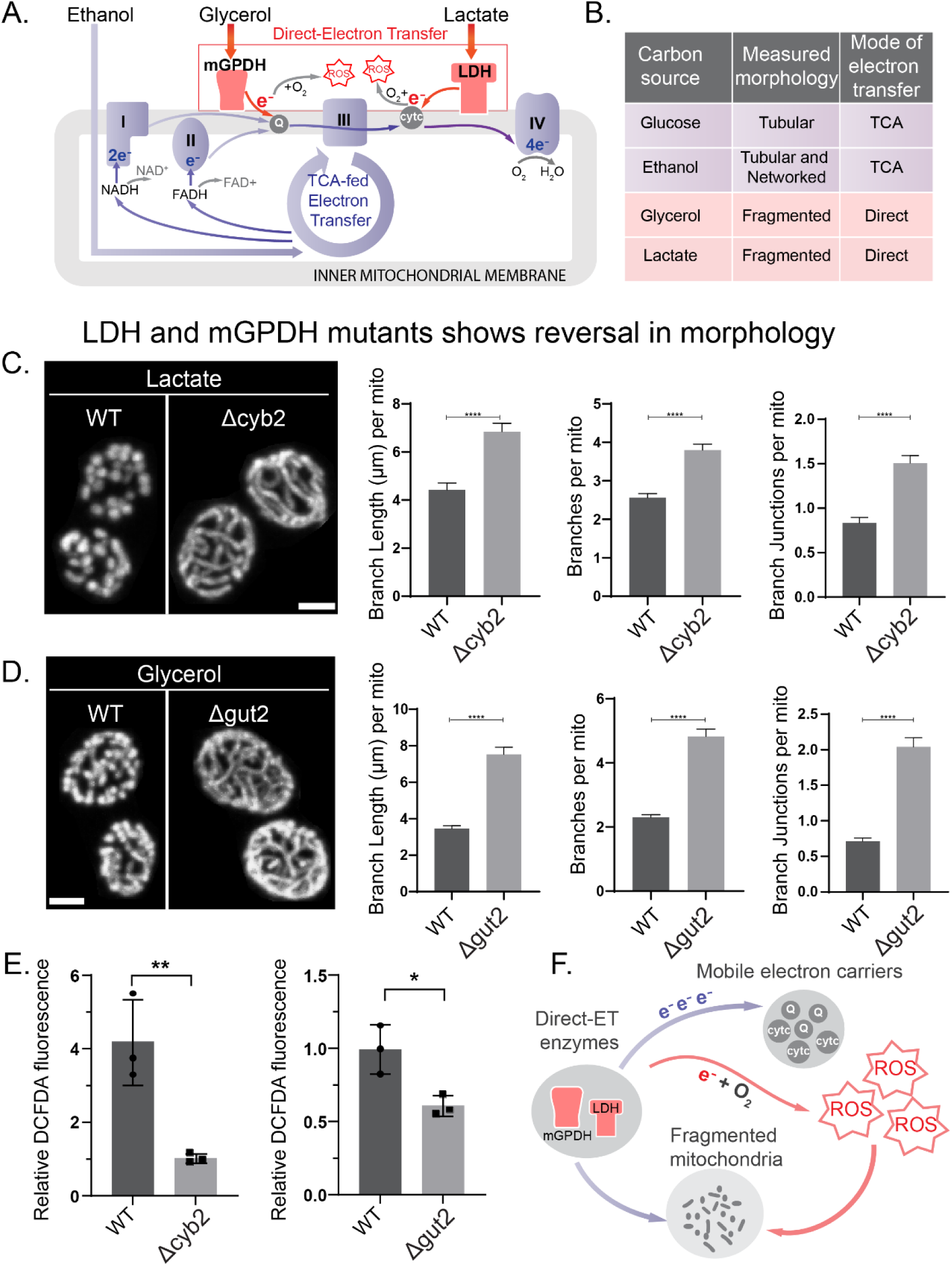
Networked mitochondria and reduced environments are restored in LDH and G3PDH mutants. (A) Schematic showing differences in electron injection into ETC when cells are grown on different carbon sources. (B) shows classification of the mode of electron entry into ETC for different carbon sources. Quantification of mitochondrial network morphology in (C) L-lactate dehydrogenase (Cyb2) deletion mutants grown in L-Lactate medium, and (D) glycerol-3-phosphate dehydrogenase (Gut2) deletion mutants grown in glycerol medium. Data represents mean ± SEM. See supplementary methods for details. (F) The redox state in Cyb2 and Gut2 deletion mutants was estimated by changes in DCFDA fluorescence intensity. Data represent mean ± SD from three biological replicates (n=3). *P < 0.05, **P < 0.01, and ***P< 0.001; n.s., non-significant difference, calculated using unpaired Student’s *t* tests.

If differences in direct electron entry determined the mitochondrial morphology, a prediction would be that preventing/reducing this direct electron injection (using mutant cells lacking the major forms of the LDH (Cyb2) or GPDH (Gut2) enzymes) should restore networked mitochondria. To test this, we first compared the mitochondrial morphologies in WT and Δ*cyb2* cells (note: see methods for growth and experiment details,) in a lactate medium. WT cells showed fragmented mitochondria, as described earlier. In contrast, Δ*cyb2* cells showed a well-networked mitochondrial morphology (Fig 4C). Next, we tested the effect of the loss of Gut2 in yeast cells in glycerol. Note: this is a challenging experiment to perform, and so the exact growth details are extensively provided in the methods section. WT cells showed the expected fragmented mitochondria. In contrast, Δ*gut2* cells showed a well-networked mitochondrial morphology (Fig 4D). If this restored network were due to reduced ROS production (from reduced direct electron transfer), the intracellular redox state in these mutants would also be expected to become more reduced. We therefore assessed intracellular redox states in these cells (as compared to WT cells). Consistently, both Δ*cyb2* and Δ*gut2* cells showed a substantially more reduced intracellular state (Fig 4E). Collectively, these data suggest that preventing direct electron entry into ETC will result in more networked mitochondria (Fig 4F).

### Altering electron flow to change ROS controls mitochondrial morphology

These data suggest that if electron flow, and thereby electron accumulation is modulated (to reduce resultant ROS), the form of the mitochondrial network could be reversibly controlled. Experimentally, this could be addressed by using a (carefully titrated) inhibition of different ETC complexes. When performed without completely ablating mitochondrial activity, this would result in blocked electron flow, alongside an accumulation/increased release of electrons. The accumulation of electrons should increase ROS. An increase in ROS (regardless of activity) should lead to fragmented mitochondria.

To test this, we selected three distinct inhibitors of electron entry into different complexes of the ETC, which function in yeast. These are: sodium azide (which inhibits complex IV), myxothiazol (which inhibits complex III), and carboxin (which inhibits complex II) (Fig 5A) (57–61). Sodium azide binds heme and copper sites in complex IV to block electron transfer from cytochrome c to complex IV. Myxothiozol blocks electron transfer from ubiquinol to the Riske-iron sulfur complex in complex III. Carboxin binds to the quinone–reduction site of complex II, thereby also preventing electron transfer (Fig 5A). Note that a careful titration of each of these inhibitors had to be first established; in order to optimize the concentrations for this experiment in prototrophic yeast cells. Extensive details are provided in the methods section and Fig. S5. In cells growing in ethanol with well-networked mitochondria, we first added sodium azide. Addition of sodium azide, within 30 min led to fragmented mitochondria, along with a concurrent drop-in mitochondrial activity (Fig 4B). To these cells, we next added a combination of antioxidants (ascorbic acid and mitoTEMPO). Within 30 min, we now observed a complete recovery and restoration of the mitochondrial network (Fig 5B). Notably, the addition of the inhibitor resulted in an increase in ROS, and this increase in intracellular oxidized state was rescued upon adding the antioxidant. However, the mitochondrial activity that reduced upon addition of sodium azide did not recover upon addition of the antioxidant (Fig 5B).

**Fig. 5.**
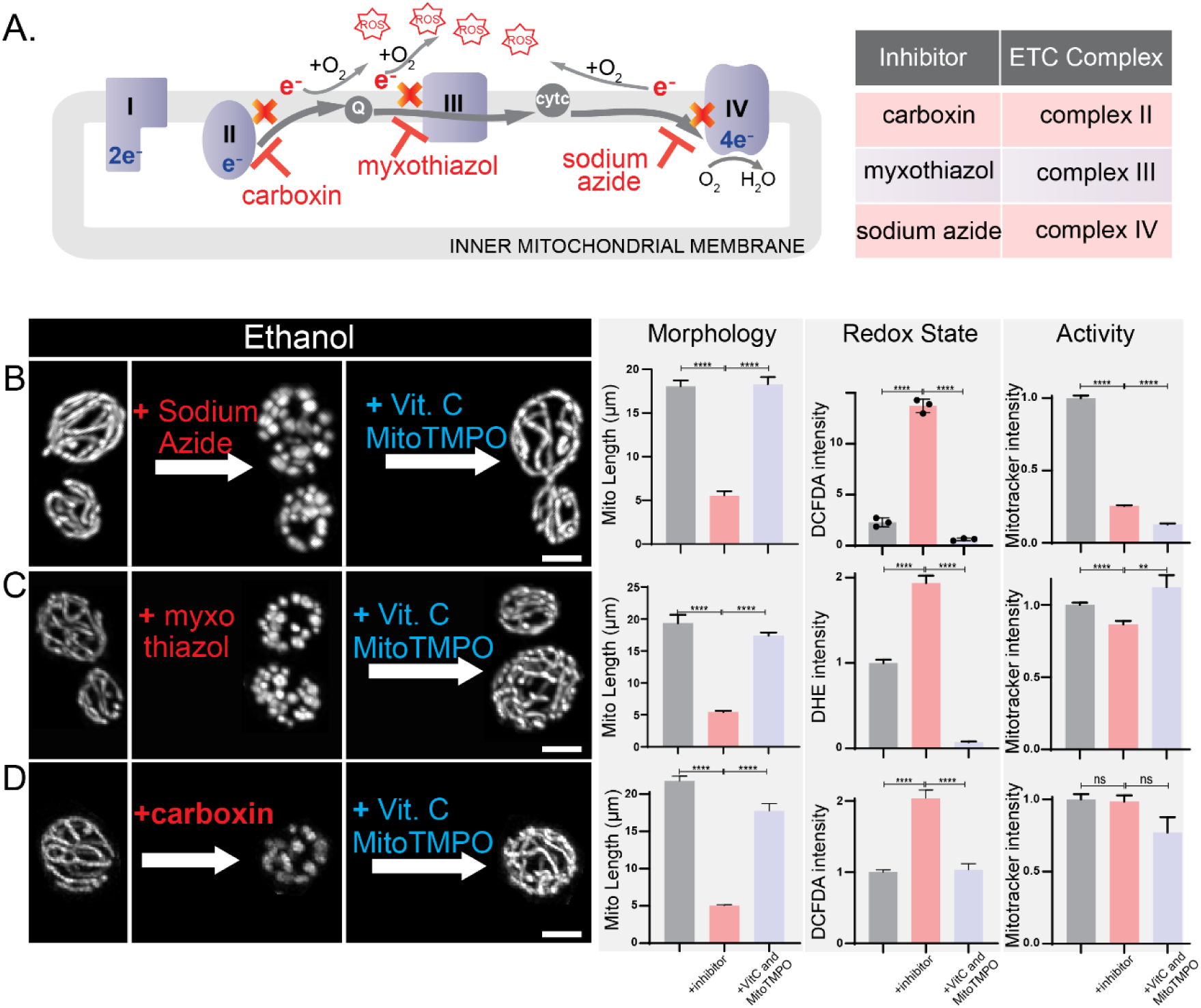
Effect of altering the nature of electron flow on mitochondrial morphology. (A, B) Schematic showing how different ETC inhibitors (classified in B) might generate ROS. (C, D, E) Changes in mitochondrial morphology for cells grown in ethanol upon addition of sodium azide (complex IV inhibitor), myxothiazol (complex III inhibitor), and carboxin (complex II inhibitor) respectively. Mitochondrial morphology is restored upon incubation of cells with an antioxidant cocktail of ascorbic acid (Vit C) and mitoTEMPO (mitochondria targeted superoxide scavenger). Mitochondrial morphology is assessed by quantifying average mitochondrial length per mitochondria, redox state is measured by evaluating changes in DCFDA or DHE fluorescence (Fig S4) and Mitotracker fluorescence is used as a proxy for measuring changes in activity. Data represents mean ±SEM. Scale bar - 2µm. *P < 0.05, **P < 0.01, and ***P< 0.001; n.s., non- significant difference, calculated using unpaired Student’s *t* tests.

We next performed similar experiments with myxothiazol (Fig 5C) and carboxin (Fig 5D), which respectively inhibit complex III and II. In each of these cases, the addition of the inhibitor resulted in a highly fragmented mitochondrial morphology (Fig 5C and 5D). Upon the addition of antioxidants, ascorbic acid and mitoTEMPO, this fragmented morphology was restored to a networked morphology. Correspondingly, upon the addition of the respective inhibitors, we also observed an increase in intracellular ROS (Fig 5C and Fig 5D), which is rescued upon the addition of the strong antioxidants. Note also that there was no correlation of mitochondrial morphology with activity in these contexts.

Summarizing, by controlling the nature of electron flow it is possible to reversibly regulate the mitochondrial network morphology. Blocking electron flow using inhibitors (of different ETC complexes) results in fragmented mitochondria and increased ROS. If this ROS is scavenged by antioxidants, the same cells will restore their mitochondrial network morphology. Notably, the changes in mitochondrial morphologies observed in the experimental conditions did not correlate with the mitochondrial activity (Fig 5B,5C & 5D).

### The dynamin-related fission GTPase Dnm1 rapidly responds to redox state changes

The mitochondrial fission and fusion machinery control the eventual mitochondrial morphology, regardless of the context. Finally, we therefore asked if the fission machinery responded to this rapid alteration of morphology by redox state. The dynamin-related GTPase Dnm1 (homologous to Drp1 in mammals) is the primary regulator of mitochondrial fission (62, 63). Dnm1 oligomerizes around and pinches off the mitochondria, and increased Dnm1 activity will result in increased mitochondrial fragmentation (63). In order to quantify fragmentation, Dnm1 activity needs to be observed *in vivo* (in the form of foci assembled on mitochondria). When native Dnm1 is tagged with a bulky fluorescent epitope (e.g. GFP) it severely affects cell growth. In order to therefore visualize native Dnm1, we engineered native Dnm1 with a short ALFA-tag at the carboxy-terminus (64), after modifying this system for use in prototrophic yeasts (Supplementary methods). These cells were engineered to stably express an ALFA nanobody (ALFA^NB^), as illustrated in the schematic in Supplementary Fig S6, and as elaborated in the Supplementary methods. Cells with native Dnm1 engineered with a short carboxy-terminal ALFA-tag behaved identically to wild-type cells, with no growth defects. Through this system, Dnm1 puncta could be unambiguously visualized in live cells, localized to the mitochondria. As Dnm1 localizes into punctae on mitochondria, this will be detected with the assembly of the fluorescent ALFA nanobody at that location (Supplementary Video S2 and Fig 6A). We next visualized and quantified the number of Dnm1 puncta in cells growing in ethanol, glycerol or lactate, where mitochondria are more networked (ethanol) or fragmented (glycerol or lactate). Cells growing in ethanol had significantly fewer Dnm1 puncta per mitochondrial length (Fig 6A and 6B) than cells in glycerol or lactate. This is consistent with lower Dnm1 activity during steady state growth in ethanol, compared to lactate or glycerol. Next, we manipulated the electron flow through the ETC, using pharmacologic treatments as in Figure 5, in order to increase ROS. We treated cells growing in ethanol with myxothiazol (as shown in Figure 5), and observed Dnm1 puncta within the same cells over time. Notably, within ∼5 minutes of adding the inhibitor, we observed a rapid, continuing increase in Dnm1 puncta formation (Fig 6C & 6D). This suggests greater Dnm1 activity in response to the oxidation associated with inhibitor addition, as a consequence of the blocked electron flow. As shown earlier, complete fragmentation of the mitochondria is observed within 20-30 min. Finally, to these same cells (with now fragmented mitochondria), antioxidants (Vit C and mitoTEMPO) were added. Within 5 minutes of antioxidant addition, the Dnm1 puncta rapidly decreased in these cells (Fig 6C & 6D), and the mitochondria become more networked (as shown earlier).

**Fig. 6.**
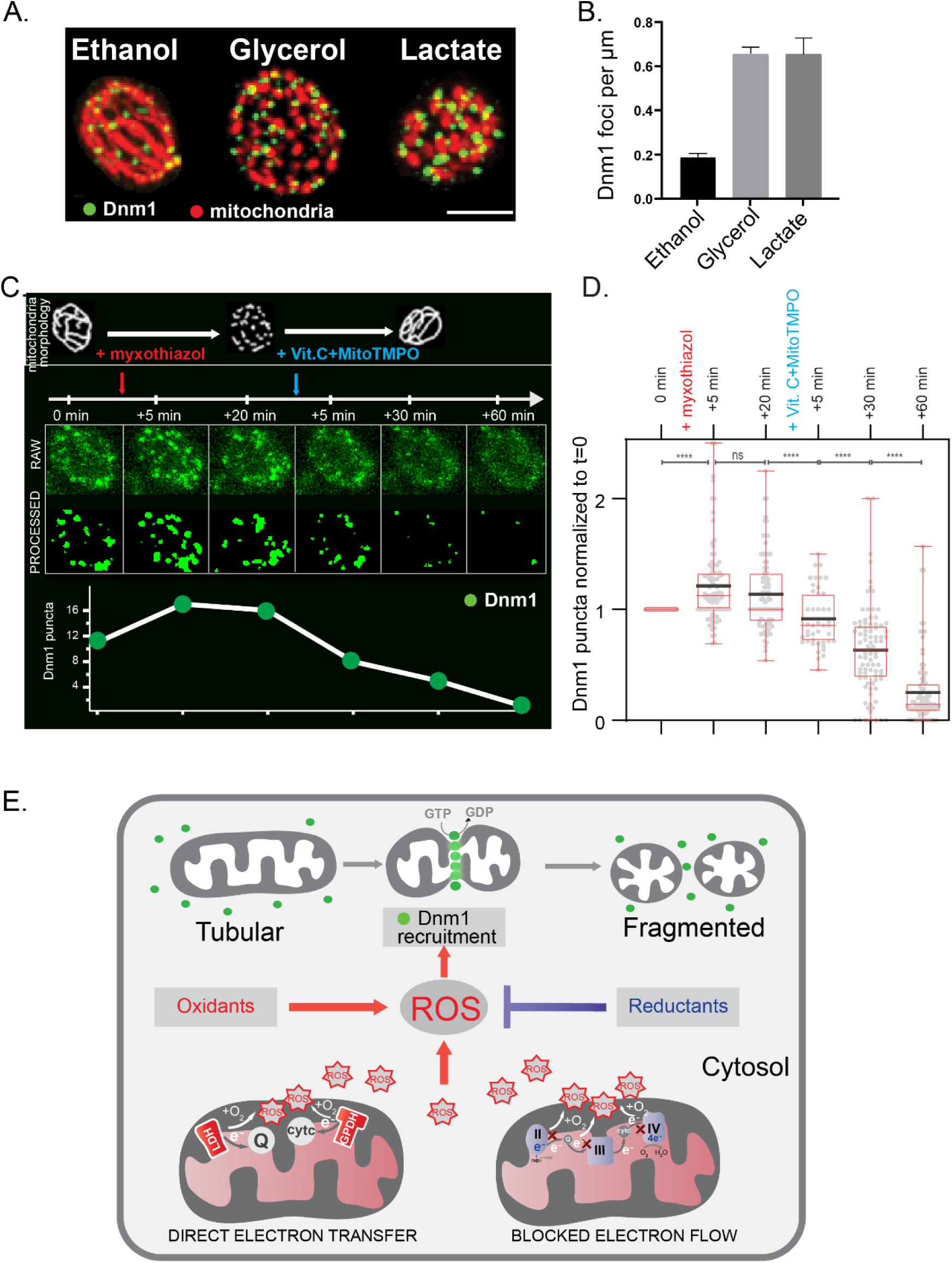
The fission GTPase Dnm1 rapidly and reversibly responds to redox state. (A) Maximum intensity projected images showing Dnm1 puncta (green) colocalized on mitochondrial tubules (red) in different carbon sources with (B) showing the quantification of the number of Dnm1 puncta in these carbon sources. Scale bar - 2µm. Cells grown in glycerol and lactate that show fragmented mitochondria also have considerably higher number of Dnm1 puncta per mitochondrial length. (C) Real-time tracking of Dnm1 puncta (green) in cells grown in ethanol. Number of Dnm1 puncta increases upon adding myxothiazol (complex III inhibitor that increases ROS and induce mitochondrial fragmentation as shown earlier) within 5 min. Upon adding an antioxidant cocktail of ascorbic acid (Vit C) and mitoTEMPO (mitochondria targeted superoxide scavenger), Dnm1 puncta reduces within ∼5 min with almost complete reduction in 30-60 min. (D) Quantification of Dnm1 puncta as a function of time (data includes time-lapse imaging for 87 cells). (E) An illustrative model proposing redox state as an integrated systems level biochemical signal to regulate mitochondrial network morphology. *P < 0.05, **P < 0.01, and ***P< 0.001; n.s., non-significant difference, calculated using unpaired Student’s *t* tests.

In summary, in contexts resulting in rapidly increased ROS, Dnm1 puncta increase within 5 minutes of the change in redox state, and the mitochondria fragment. This process is rapidly reversible. Upon adding antioxidants, the Dnm1 puncta decrease immediately and the mitochondria become more networked. Collectively, these data reveal that the activity of the central fission machinery (Dnm1) rapidly responds to changes in ROS, to actively remodel the mitochondrial network in tune with the redox state of the cell.

## Discussion

Mitochondria are dynamic organelles that transition between networked and fragmented forms, and it has remained unclear how this relates to mitochondrial activity. Here, we find that mitochondrial form is directly controlled by the intracellular redox state, which is in turn regulated by the nature of metabolic processes in the cell that alter ROS. A synthesis of these results is summarized in Fig 6E, suggesting a chemical basis of how mitochondrial form is regulated based on intracellular redox state. The intracellular redox state is determined by ROS sources, primarily produced by the mitochondria itself - but which can also come from other sources. The amount of ROS produced will depend on the nature of metabolism within the cell. The nature of electron entry into the ETC, and therefore electron flow through the ETC is not unidimensional. In some metabolic environments, there can be increased direct electron entry/injection into the ETC, as occurs with the yeast LDH or GPDH enzymes during growth in lactate or glycerol. Whenever this occurs, there is high ROS generation as a consequence, leading to an oxidized intracellular environment which results in fragmented mitochondria. If direct electron entry into the ETC is curtailed (through either genetic or pharmacologic approaches), there is reduced ROS, and a more networked- mitochondria. This explains why cells growing in different carbon sources all of which require mitochondrial respiration, can show distinct mitochondrial network morphologies. In contexts requiring high direct electron transfer that leads to ROS generation (e.g. lactate, glycerol), cells have fragmented mitochondria. Importantly, this ROS-dependent control of mitochondrial form is rapidly reversed (in minutes) by altering only the redox state. For example, reducing the intracellular environment in cells growing in lactate will result in networked mitochondria, or adding oxidants will fragment mitochondria in cells growing in ethanol. Additionally, this network can also thereby be independent of activity (in terms of mitochondrial oxygen consumption or respiration). Mechanistically, the mitochondrial fission GTPase Dnm1 responds rapidly to changes in cellular redox state. Dnm1 puncta rapidly increase upon an increase in ROS, along with mitochondrial fragmentation. Correspondingly, strong antioxidants can reverse this Dnm1 puncta formation along with fragmentation. We thus speculate that the rapid change in redox state (through changes in ROS) signal to control fission activity, to thereby reversibly control mitochondrial morphology independent of mitochondrial activity itself.

Electron entry into the ETC (and therefore resultant electron flow) can occur and different locations, and illustrates an underappreciated versatility of the ETC machinery. In a ‘text-book’ scenario where substrates fuel the TCA cycle, the TCA cycle generates reducing equivalents that supply the electrons to drive the ETC. In these cases, electron flow is tightly controlled and with minimal leakage of electrons (and therefore minimal reactive oxygen species generated). In contrast, when cells grow on carbon sources where their metabolism is coupled with a ubiquinone or cytochrome c, electrons enter ETC through direct electron injection (65), leading to increased ROS. In such contexts, the mitochondria will be fragmented, independent of mitochondrial ‘activity’. While mechanisms of direct electron injection/entry remain poorly understood, there is considerable evidence of its common occurrence in different growth conditions, and is observed ubiquitously across species. There are enzymes that transfer electrons directly to ubiquinone pool including proline dehydrogenase (PRODH) involved in proline catabolism, mitochondrial GPDH or GPD2, dihydro-orotate dehydrogenase (DHOODH) associated with pyrimidine synthesis, electron transfer flavoprotein:ubiquinone oxidoreductase (ETF: QO) that transfers electrons from beta-oxidation of fats to ubiquinone (53, 66). In all these contexts, cells appear to have high ROS generation (67, 65). Why might such ‘direct’ electron entry mechanisms exist in the mitochondria? A speculative argument would be that this increases the versatility of mitochondrial functions in order to enable cells to utilize a range of carbon sources that would otherwise not easily be metabolized. In contexts where the activity of TCA cycle is reduced, electron acceptors such as ubiquinone or cytochrome c can receive electrons from other substrates that can be oxidized easily. This would allow the ETC to continue to function, and sustain the energetic and redox needs of the cell. This would also result in metabolically plastic cells that are capable of growing in diverse carbon/nutrient environments, as opposed to specialized cells that grow in a limited carbon source.

In the context of mitochondrial morphology, any mechanism therefore that increases the oxidative environment of the cytosol can lead to the fragmentation of mitochondria. This means exogenous generation of ROS through oxidizing agents, or atypical oxidation of fatty acids (68) or unchecked peroxisomal activity (69) could all result in a fragmented mitochondrial morphology. Such an interpretation can now explain observations from mutants or disease conditions where fragmented mitochondria are observed – without a change in mitochondrial activity (15, 16, 29). Here, it is likely that the intracellular state is more oxidized due to perturbed electron flow from direct electron injection, and/or exogenous ROS generating species. Finally, mutations or inhibition of ETC complexes that block electron flow would also result in an accumulation of electrons, leading to ROS generation through one-electron reduction of molecular oxygen to superoxide, or indirectly by facilitating reverse electron transfer, leading to the fragmentation of the mitochondria (as we exploit in this study). In these contexts, the activity of mitochondria might be compromised (e.g. complex IV inhibition using sodium azide) or remain unchanged (e.g. complex II inhibition using carboxin). Regardless, all these conditions will result in fragmented mitochondria due to increased intracellular oxidation. This therefore need not be limited to energy stress alone, which is one form of stress that can fragment the mitochondria (70). In any of these conditions, if the intracellular environment can be reduced or ROS removed (for example by using strong antioxidants) the mitochondrial morphology will restore to a networked form. This can be independent of mitochondrial activity itself – for example the action of ETC inhibitors will reduce mitochondrial activity (and fragment the mitochondria), but adding antioxidants will now allow the network to reestablish. Our study also highlights the value of using a clean, reductionist approach to address this question. Prototrophic yeast cells are metabolically versatile, and metabolize different carbon sources – all of which requires mitochondrial respiration. Using these defined drop-out/add back conditions where we could independently assess mitochondrial activity, potential and morphology allowed a precise correlation of what controls form to be established. In a metabolically specialized cell, it would be impossible to dissect out how mitochondrial form might be regulated by direct electron entry vs other forms of electron entry that are supported by reducing equivalents derived from the TCA cycle, due to distinct metabolic outputs.

Cells can rapidly control form based on the redox state of the cell, by coupling the ROS response to the activity of the fission machinery. The fission GTPase Dnm1 responds almost immediately to changes in the redox state of the cell, increasing in activity (as observed with more Dnm1 puncta) when electron flow is altered (which leads to increased ROS generation). This phenomenon is also rapidly reversible. A prediction emerging from this study is therefore the likely presence of signaling mechanisms that couple redox state changes to regulated mitochondrial fission through Dnm1 activity. Such a system to communicate with effectors of final mitochondrial form should regulate of the relative dynamics of fission vs fusion, alongside other factors such as the recruitment of cytoskeletal determinants of mitochondrial motility. Interestingly, recent studies also reveal distinct types of fission in cells, relating to degradation vs biogenesis (71). Our study therefore reconciles multiple studies that identify such mechanisms as regulators of mitochondrial form (72, 73, 16, 74, 70), while delineating a path for future inquiry to identify upstream regulators of mitochondrial form based on cellular redox state changes as a function of metabolic state.

## Materials and Methods

The specifics of the media and growth conditions for different cells, mitochondrial imaging and analysis, mitochondrial activity and redox assays, and metabolic analysis are detailed in the Supplementary Materials and Methods section of the SI Appendix.

### Cell culture and biochemical assays

A prototrophic CEN.PK strain of *Saccharomyces cerevisiae* was used in all the experiments. Gene deletions, chromosomal C-terminal tagged strains were generated by PCR mediated gene deletion/tagging (75). Mitochondria targeted mNeon strain (Mito-mNeon green) is described in (76). Media compositions and specific growth conditions are described in the Supplementary Materials and Methods section of the SI Appendix. The fluorescence intensity for plate reader based biochemical assays was measured using Thermo Varioskan LUX multimode plate reader. Detailed protocol is described in SI Appendix. The fluorescence intensities for each sample were normalized using OD_600_ of that sample and relative fluorescence intensity was calculated and plotted.

### Protein extraction and Western blotting

Total protein was extracted using trichloroacetic acid (TCA) precipitation and blots were quantified using FIJI software. Detailed protocol is described in SI Appendix.

### Basal OCR measurement

Agilent Seahorse XFe24 analyzer was used for basal OCR measurements. The detailed methods are described in SI Appendix.

### Estimation of GSH levels by LC-MS/MS

The steady state levels of GSH were estimated by quantitative LC-MS/MS methods as described previously (77). Detailed methodology is described in the SI Appendix and peak area measurements are listed in the supplement file Dataset S1.

### Fluorescence Imaging and Analysis

Fluorescence imaging experiments were performed on live cells on an inverted laser scanning confocal microscope. For all quantifications of mitochondrial morphology, cells containing mneonGreen tagged mitochondria were used. Mitochondrial membrane potential was measured by staining cells with Mitotracker CMXRos. Redox state was measured in live cells by evaluating changes in DCFDA or DHE fluorescence. Custom- written macros and plugins in ImageJ were used for image analysis. Please refer to SI Appendix for detailed imaging pipeline.

## Acknowledgments

We acknowledge extensive use of the Central Imaging and Flow Facilities, as well as the mass spectrometry facility in the NCBS and inStem campus. We acknowledge Ssaumya Seth for help in generating the alpha-tag Dnm1 strain. We acknowledge Suresh Subramani, Jitu Mayor, MK Mathew, Amit Singh, Christian Frezza, Richa Ricky and members of the SL lab for feedback at various stages of this study. GS acknowledges a Research Associate Postdoctoral fellowship (DBT-RA) from the Dept. of Biotechnology, Govt. of India. SL acknowledges funding support from DBT-Wellcome Trust India Alliance Senior Fellowship (IA/S/21/2/505922), and S. Ramachandran National Bioscience Award for Career Development from the Dept. of Biotechnology, Govt. of India.

