## Supplemental File for "Active control of mitochondrial network morphology by metabolism driven redox state"

**This PDF file includes:**

- Supporting text
- Figures S1 to S7
- Legends for Movies S1 to S2
- Legends for Datasets S1
- SI References

**Other supporting materials for this manuscript include the following:**

- Movies S1 to S2
- Datasets S1
- Software S1

### Material and methods

#### Media and growth conditions

The prototrophic CEN.PK strain of *Saccharomyces cerevisiae* was used in all the experiments. For experiments involving mitochondrial imaging and morphology analysis, mitochondria-targeted mNeonGreen expressing strain (mt-mNG) was used (1). Cells were grown in a synthetic complete medium (0.67% yeast nitrogen base with ammonium sulfate, supplemented with 2 mM amino acids) with five different carbon sources - 2% glucose (glucose medium), 2% glycerol (glycerol medium), 1% glycerol-2% ethanol (glycerol-ethanol medium), 2% ethanol (ethanol medium) and 2% DL-lactate (lactate medium). For experiments involving the L-lactate dehydrogenase deletion mutant (cyb2), 2% DL-lactate or 2% L-Lactate was used as indicated. For experiments involving mitochondrial glycerol-3-phosphate dehydrogenase (gut2) mutant, cells were not viable in glycerol media therefore we first grew cells in glycerol-ethanol medium for 2 hours at a starting OD600~ 0.3 and then shifted these cells to glycerol medium for 4 hours. Mitotracker CMXRos (catalog no M7512), H2DCFDA (catalog no C6827) and DHE (catalog no D11347) were purchased from Thermo Fisher Scientific and were added to a final concentration of 200 nM, 10 µM and 5 µM respectively in cells. Cells stained with these dyes were incubated at 30°C in a shaking incubator for 30 mins and then were washed and resuspended in fresh media to be used for microscopic imaging or plate reader-based assays.

#### Generation of ALFA-tagged Dnm1 strain

The pABY-c15 plasmid (2) that express nanobody against ALFA tag (NbALFA) linked to mNeonGreen under TEF promoter was cloned into a yeast integrating vector pCfB2513 (pCfB2513 was a gift from Irina Borodina, Addgene plasmid #67543 (3)) by Gibson assembly cloning. The TEF promoter was replaced by G6PD promoter by Gibson assembly to generate G6PD-NbALFA-mNeonGreen with a hygromycin resistance selectable marker, flanked by X-2 chromosomal integration sequence. The region of interest was PCR amplified and transformed into yeast cells to generate stable, chromosomally integrated Nb ALFA-mNeonGreen expressing cells. The transformants were selected using Hygromycin resistance and the integration was confirmed by colony PCR. In this strain expressing the NbALFA-mNeonGreen, Dnm1 was C-terminally tagged using ALFA tag by PCR mediated gene tagging strategy (4) using the pYM17-ALFA-natNT2 plasmid (2). The transformants were selected using NAT resistance and C-terminal tagging was confirmed by colony PCR. Fig S6 depicts the workflow for strain construction.

#### Plate reader assay for Mitotracker, H2DCFDA and DHE

Cells were grown to $OD_{600}$~0.6, 1 $OD_{600}$ cells were collected by centrifugation (1000 x g, 1 min at room temperature (RT)), and resuspended in 1 ml fresh media. Mitotracker CMXRos or H2DCFDA were added to a final concentration of 200 nM or 10 µM respectively and incubated at 30°C in a shaking incubator for 30 mins. Cells were washed and resuspended in 1 ml fresh media in case of Mitotracker CMXRos and 500 µl in case of H2DCFDA. 300 µl of this was aliquoted into 96 well plates in replicates. Fluorescence was measured using Thermo Varioskan LUX multimode plate reader at 579/599 excitation/emission wavelengths for Mitotracker CMXRos and 495/520 for H2DCFDA  Fluorescence readings were normalized using $OD_{600}$ of each sample and the relative fluorescence intensity was calculated. Statistical significance was calculated using unpaired Student's t-test (GraphPad prism 9.0.1).

#### Mitochondrial inhibitors, reducing and oxidizing agents

Cells were grown in indicated media to an $OD_{600}$~ 0.6 and 1 $OD_{600}$ cells were collected by centrifugation (1000 x g, 1 min at RT) and resuspended in 1 ml fresh media. For experiments involving $H_2O_2$ treatment, cells were treated with $H_2O_2$ at a final concentration of 5mM for 30 minutes. For experiments involving GSH treatment, cells were treated with GSH at a final concentration of 20 mM for 30 minutes. For experiments involving GSH addition to $H_2O_2$ treated cells, cells were incubated with $H_2O_2$ for 30 minutes followed by incubation with GSH at a concentration of 20 mM for another 30 minutes at 30°C in a shaking incubator. For oxidative stress rescue using ascorbate (Vit C) - mitoTEMPO cocktail, cells treated with mitochondrial inhibitors (sodium azide - 500 µM, myxothiazol -20/100 nM, carboxin - 2mM, 20 mins incubation) were washed and incubated with 100 mM Vit C and 1 mM mitoTEMPO for 30 or 60 minutes, at 30°C in a shaking incubator. Cells were then washed and resuspended in fresh media to be used either microscopic imaging or plate reader-based assays.

**Protein extraction and Western blotting**

Cells were grown in appropriate media to $OD_{600}$~ 0.6 and centrifuged (1000 x g, 2 min at RT). Total protein was precipitated using trichloroacetic acid (TCA), resuspended in SDS/glycerol buffer by heating at 90°C for 10 minutes and centrifuged. The supernatant was collected and total proteins were estimated by BCA protein estimation assay (BCA assay kit, G-Biosciences). The samples were normalized to ensure the equal protein amounts in all the samples. The protein samples were run on 4–12% precast bis-tris gels (Invitrogen, NP0336BOX), using MOPS running buffer (50 mM tris, 50 mM MOPS, 0.1% SDS, and 1 mM EDTA). The relevant portion of the gel is transferred to a nitrocellulose membrane (GE Healthcare, 10600003), while a different region was stained with coomassie blue for loading control. The blots were developed using the following antibodies: anti-HA mouse (Sigma-Aldrich 11583816001), anti-FLAG mouse (Sigma-Aldrich F1804), anti-Cox2 mouse (Invitrogen MTCO2 459150). HRP-conjugated secondary antibodies from Sigma-Aldrich (mouse and rabbit) were used. Chemiluminescence was detected by using Western Bright ECL HRP substrate (Advansta, K12045).

**Maleimide-PEG assay**

Cells were grown in appropriate media, 10 $OD_{600}$ cells were collected and ice-cold trichloroacetic acid (TCA) was added to cells at a final concentration of 10%, followed by 15 mins incubation on ice. Cells were centrifuged at 1500 x g, for 2 mins at 4°C and proteins were precipitated using TCA as described above. The protein pellet was washed with 1ml of ice-cold 100% acetone and centrifuged at 20000 x g for 5 mins at 4°C. The pellet was dried using a speed-vac at RT for 2-5 mins and resuspended in 300 µl Tris-SDS buffer (100mM Tris (pH 8), 1%SDS, 1mM EDTA (pH 8), 1mM PMSF). 50 mM mPEG was prepared in Tris-SDS buffer and the reaction mixture was set up by adding 50 µl protein sample, 2.4 µl mPEG and 7.6 µl Tris-SDS buffer. For control samples without mPEG, 50 µl protein sample and 10 µl Tris-SDS buffer was mixed. The reaction mixtures were incubated at 37°C for 1 hour in a thermomixer and the reaction was stopped by adding 20 µl 4 x SDS-sample buffer. The samples were heated at 95°C for 5 mins, run on 4–12% precast bis-tris gel and western blot was done using anti-FLAG antibody. For quantifying the reduced proteins, the intensity of the slow migrating bands in the mPEG-treated samples were calculated and normalized to the band intensity of mPEG non-treated control for the same sample (which corresponds to the total protein levels). The relative reduced protein levels were plotted.

**Seahorse assay**

The sensor cartridge plate was hydrated overnight using the XF-calibrant solution as per the manufacturer's instructions. The sensor cartridge was equilibrated in the Seahorse XFe24 analyzer prior to the start of the assay. The culture plate was coated with poly-L-lysine (50 µl per well), followed by incubation for 1 hour at RT. Excess poly-L-lysine was then removed and the plate was dried for 30 minutes at 30°C. The cells were grown in appropriate media to an $OD_{600}$~ 0.6. The samples were aliquoted in each well such that the final cell number in each well was ~$3x10^5$. The plate was centrifuged for 2 mins (100xg, acceleration 2, brake 2) and incubated at 30°C for 30 minutes. The plate was loaded to the Seahorse XFe24 analyzer and basal OCR measured over time. Three measurements were taken with intermittent 2 min mixing and waiting steps. The basal OCR reading in each well was normalized to the $OD_{600}$ values for cells in the corresponding well.

**GSH measurement by LC-MS/MS**

For measuring reduced glutathione (GSH) levels, cells were grown in appropriate media to an $OD_{600}$~ 0.6. The metabolites were extracted by collecting 5 $OD_{600}$ cells, quenched in 60% methanol at -40°C as described in Walvekar et al (5). The extracted metabolites were separated on Waters Acquity UPLC system (using a Synergi 4µm Fusion-RP 80 Å LC column (150 x 4.6 mm, Phenomenex), and measured as described earlier (Walvekar et al., 2018), in positive polarity mode. Solvents used:0.1% formic acid in water (solvent A) and 0.1% formic acid in methanol (solvent B). ABSciex QTRAP 6500 mass spectrometer was used, data acquisition was done using Analyst 1.6.2 (Sciex), and data analysis was done using MultiQuant version 3.0.1. Detailed flow parameters are described elsewhere (Walvekar et al., 2018). The parent and product ion masses used for the analysis are listed in supplement file 1. The peak areas for GSH were calculated and values were plotted relative to glucose. The retention time for GSH and peak area obtained after analysis are listed in Dataset file S1. The statistical significance was calculated using unpaired Student's t-test (GraphPad prism 9.0.1).
*Note: For GSH level estimation, samples were run on LC-MS/MS immediately after extraction since storage of extracted samples resulted in decrease in GSH peak areas over time, possibly due to oxidation.*

**Fluorescence microscopy and Image analysis**

All Fluorescence imaging experiments were performed on living cells using an Olympus FV3000 or Zeiss LSM 980/780 inverted confocal laser scanning microscope. A 60X or 63X oil objective (NA ~1.4) was used for focusing the excitation light and fluorescence was collected using GaAsP-PMT based high-sensitivity detectors. We used a 488 nm laser for exciting cells tagged with mitochondria targeted mNeonGreen as well as cells stained with H2DCFDA and DHE. Mitotracker-red was excited using 561 nm laser excitation. Cells were imaged on custom made 35 mm imaging dishes with an acid cleaned coverslip (using 2:1 ratio of nitric acid and hydrochloric acid (6) for 2 hours) attached to the bottom. Using customized dishes with acid-cleaned high-quality coverslips (Thermo Scientific™ Gold Seal™ No #1) allowed us to acquire high quality 3D confocal z-stacks of mitochondria with minimal background. Yeast cells being non-adherent were immobilized using custom-designed agarose pads (made using 1% ultrapure agarose in distilled water). For quantification of mitochondrial morphology in different conditions, 3D confocal images with an x,y scaling of ~50 nm per pixel and z-scaling of ~0.4 µm per slice were acquired as per the optimal Nyquist sampling parameters. For intensity measurements, a larger field of view was used to excite and collect fluorescence from a large population of cells.

Fig. S1A shows the basic image analysis pipeline. First, raw 3D image stacks from our confocal microscope were deconvolved using Huygens Professional software 17.10 (SVI). Fig. S1B shows the comparison of axial section of $xz$ and $yz$ planes of raw and deconvolved images; highlighting significant reduction in axial stretching of objects for deconvolved images. We also tested DeconvolutionLab2 (2), an ImageJ plugin, in conjunction with a synthetic psf generated by PSF generator (3) for deconvolving fluorescence images using Richardson-Lucy algorithm (20 iterations) that also gave us comparable improvement in spatial resolution as compared to Huygens package. Deconvolved image stacks are then processed in ImageJ by either available plugins or custom written routines. Unlike mammalian cells that are flat and have most of their mitochondria in a single plane, yeast cells are round and relatively smaller in size with mitochondria spread throughout the cellular volume therefore we need to do image thresholding and quantification on 3D image-stack to capture the true state of mitochondrial networks in yeast cells. Fig. S1C highlights how image quantification using a 2D maximum intensity image can lead to incorrect assessment of mitochondrial morphology. For quantifying mitochondrial morphology in 3D z-stacks, we used an ImageJ plugin Mitochondria analyzer (7) that has been shown (8) to perform robust structural quantification of mitochondrial network morphology as compared to other methods for both 2D as well 3D fluorescence image stacks. For quantification of fluorescence intensity in case of dye-stained cells a custom script was written in ImageJ (software file S1) to allow quantification of pixels that had signal intensity above the background threshold and to reject pixels that were hot (saturated). This was important as cells in different metabolic, chemical and genetic backgrounds showed a wide variation in fluorescence intensities therefore we needed an automated pipeline of image quantification across all treatments. For fluorescence quantification using microscopic images, each data point represents average fluorescence intensity over a field of view of 20-50 cells; enough field of views are taken for each condition in order to have at least ~150-300 cells for each treatment.

**Dnm1 imaging and analysis**

To visualize Dnm1 localization in yeast cells engineered with the ALFA-tag system, the following experimental pipeline was employed. Yeast cells grown in ethanol were added to poly-lysine coated dishes. A suitable field of view (FOV) containing the cells was selected and tracked in real-time. To avoid photobleaching, confocal z-stacks were acquired at sparse time intervals (as indicated in Fig. 6C & 6D) both before and after the addition of myxothiazol and antioxidants (see Fig S7 for comparison of imaged regions with unexposed regions at 60 min timepoint). Sample preparation, including the concentration of myxothiazol and antioxidants, and image acquisition was performed as described in the methods above.

To analyze and quantify Dnm1 punctae in cells, a custom imaging pipeline was set up in ImageJ which involved the following steps: (1) **Pre-processing**: This involved creating an aligned stack of maximum intensity projected images for all time-points for each FOV. This was done using Linear Stack Alignment with SIFT multichannel plugin in FIJI package of ImageJ. (2) **Segmenting punctae**: Aligned time-resolved stacks were further processed using a clustering algorithm to identify and segment the localized Dnm1 structures. A custom script was written in ImageJ using the the k-means Clustering plugin (IJ plugins toolkit) to implement this step. (3) **Quantifying punctae distribution:** Regions of interest (ROIs) were manually drawn in segmented image stacks to choose cells that could be located and tracked in all time-frames. For analyzing these ROIs, a custom ImageJ macro was written to quantify the number of Dnm1 punctae in each cell as a function of time.

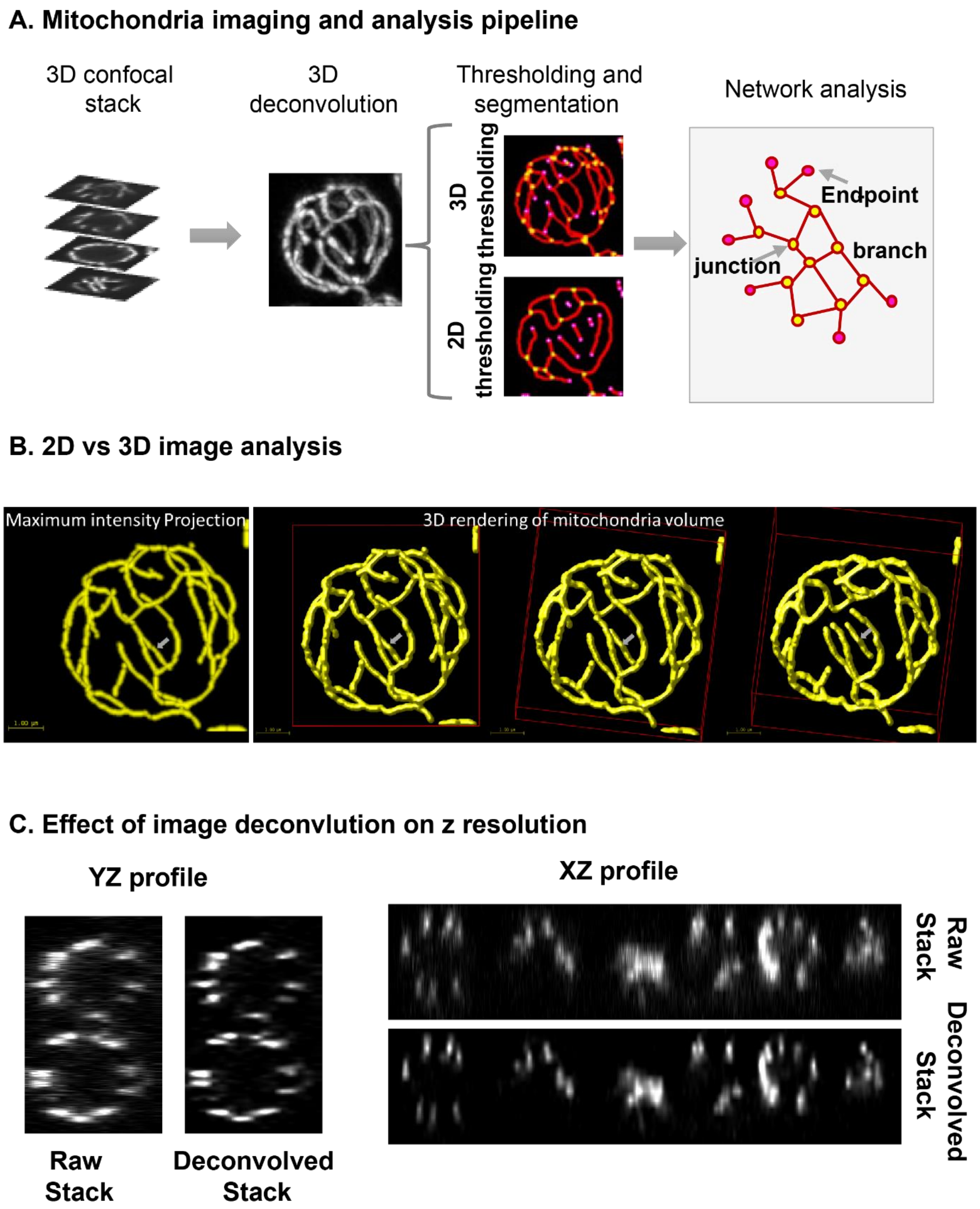
A. Mitochondria imaging and analysis pipeline
3D confocal stack
3D deconvolution
Thresholding and segmentation
Network analysis
3D thresholding
2D thresholding
Endpoint
branch
junction
B. 2D vs 3D image analysis
Maximum intensity Projection
3D rendering of mitochondria volume
C. Effect of image deconvlution on z resolution
YZ profile
XZ profile
Raw Stack
Deconvolved Stack
Raw Stack
Deconvolved Stack

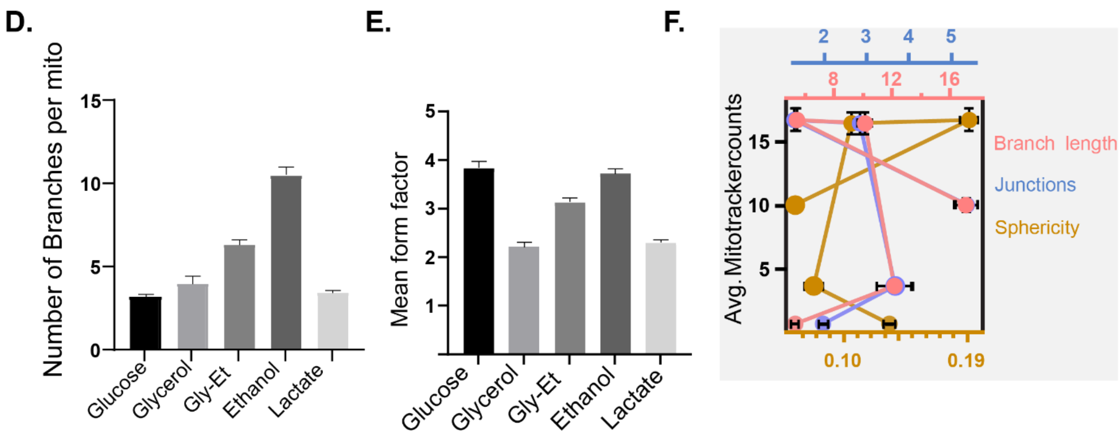


**Fig. S1.** (A) Basic workflow for mitochondrial imaging. (B) Example illustrating use of 2D vs 3D image analysis of mitochondrial network. Image on the left is a 2D maximum intensity projection (MIP) of the z-stack whereas the image-panel on the right represents 3D renderings of the same z-stack at different viewing angles. Grey arrow highlights the mitochondrion that in 2D MIP image appears as part of a single network, however 3D rendering of the same image shows that the highlighted mitochondria is in fact a short fragment thereby highlighting the need to use the 3D z-stack for image quantification instead of 2D projections of z-stack. (C) Effect of image deconvolution on axial stretching of objects. (D, E) Quantification of number of branches per mitochondria and mean form factor for different media conditions. (F) Plot of mitochondrial activity (mitotracker counts) versus various morphological parameters showing lack of correlation between mitochondrial activity and network morphology.

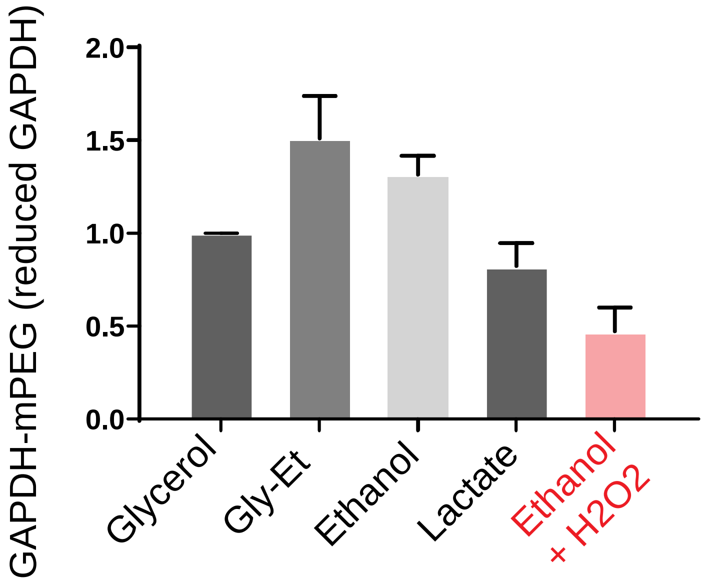


**Fig. S2.** Cytosolic oxidative state in different respiratory carbon sources as measured by electrophoretic mobility shift of mPEG bound to cytosolic GAPDH enzyme, in a western blot (using anti-FLAG antibody). As a control, we supplemented ethanol-grown cells with hydrogen-peroxide. Data represents mean ± SD from two biological replicates (n=2).

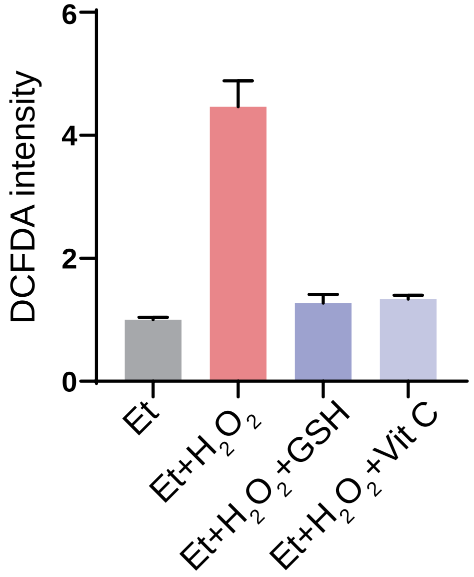


**Fig. S3.** Measuring changes in cellular redox state by using DCFDA fluorescence when ascorbate (Vit C) instead of GSH is added to cells. For ease of representation, Vit C data is added as an additional column to data already shown in Fig 3E (middle panel). Data represents mean ±SEM (see material and methods for details).

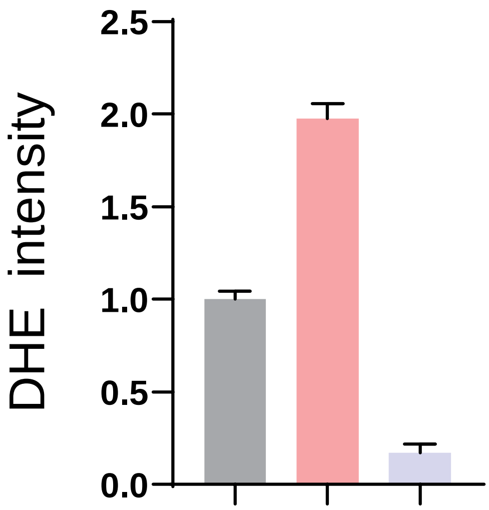


**Fig S4.** Measuring changes in cellular redox state using DHE fluorescence when complex II inhibitor carboxin is added to cells growing in ethanol. The experimental conditions are the same as those used for data shown in Fig 5D where DCFDA is used as a redox indicator. Data represents mean ±SEM (see material and methods for details).

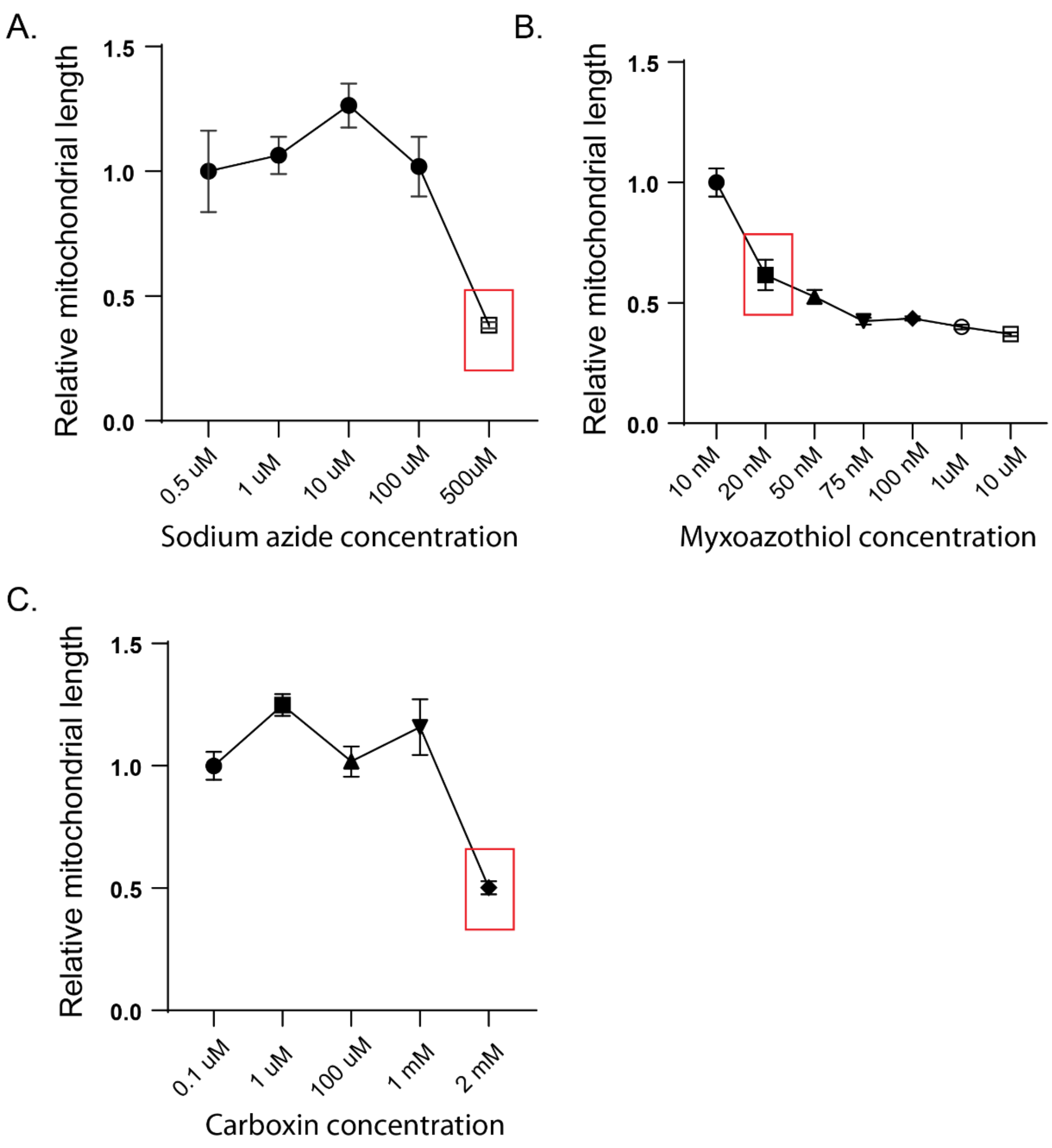


**Fig. S5.** Data showing changes in mitochondrial morphology as a function of different concentrations of mitochondrial inhibitors, (A) sodium azide, (B) myxoazothiol and (C) carboxin. Total branch length per unit mitochondria was used as a proxy for representing changes in mitochondrial morphology; shorter branch lengths would indicate more fragmented mitochondria. A careful titration of mitochondrial inhibitors was necessary to find the minimum inhibitor concentration that is sufficient to cause visible mitochondrial fragmentation in most cells but may not be high enough to completely inhibit mitochondrial activity. This is different than inhibitor concentrations generally reported in the literature where the aim is to completely inhibit or kill mitochondrial activity. This minimum inhibitor concentration found for each inhibitor is highlighted by the red rectangle in the figure.

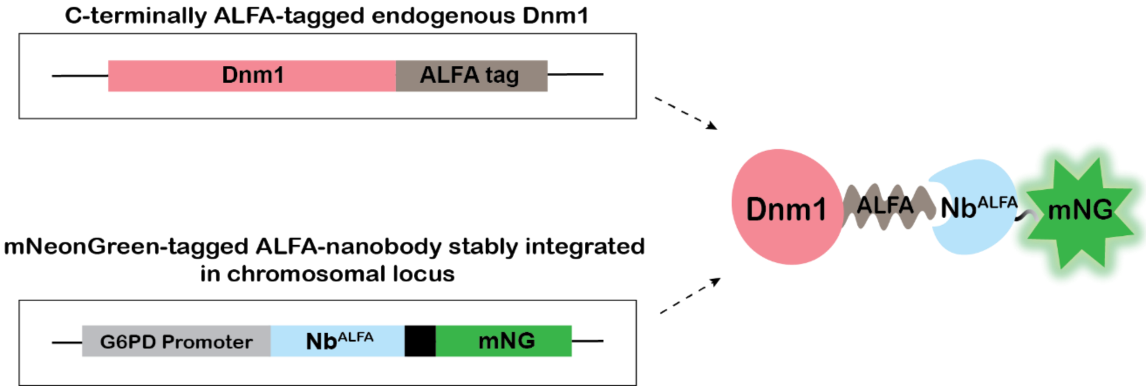


**Fig. S6.** Schematic indicating generation of Dnm1 strain with ALFA-tag system.

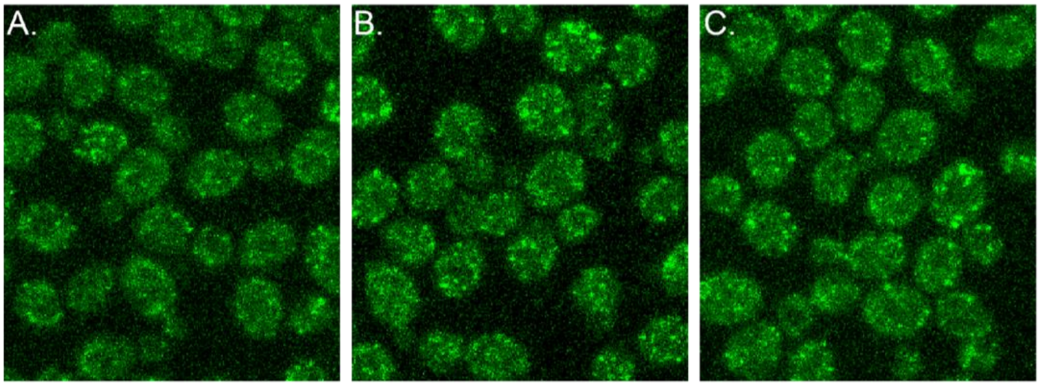


**Fig. S7.** (A) Cells showing dramatic reduction of Dnm1 punctae at 60 mins post addition of ascorbate (Vit C) and mitoTEMPO to cells that were treated with myxoazothiol. To rule out the effect of photobleaching, adjacent regions of cells (not exposed to any excitation light) indicated by (B) and (C) were also imaged. These unexposed regions also show a dramatic loss of Dnm1 punctae upon addition of antioxidants as shown in (B) and (C).

**Movie S1 (separate file).** Time-lapse images showing differences in mitochondrial network morphology between cells grown in ethanol vs lactate.

**Movie S2 (separate file).** Time-lapse images of cells showing Dnm1 punctae (green) colocalized on mitochondrial tubules (red) in different carbon sources. Mitochondria were visualized with Mitotracker Red in an ALFA-tagged Dnm1 strain (see material and methods for details).

**Dataset S1 (separate file).** Microsoft excel file containing retention times and peak areas for GSH measurement by LC-MS/MS in different carbon sources.
